# Identification and Elimination of Antifungal Tolerance in *Candida auris*

**DOI:** 10.1101/2022.12.22.521687

**Authors:** Samira Rasouli Koohi, Shamanth A. Shankarnarayan, Clare Maristela Galon, Daniel A. Charlebois

## Abstract

Antimicrobial resistance is a global health crisis to which pathogenic fungi make a substantial contribution. The human fungal pathogen *C. auris* is of particular concern due to its rapid spread across the world and its evolution of multidrug resistance. Fluconazole failure in *C. auris* has been recently attributed to antifungal “tolerance”. Tolerance is a phenomenon whereby a slow growing subpopulation of tolerant cells, which are genetically identical to susceptible cells, emerges during drug treatment. We use microbroth dilution and disk diffusion assays together with image analysis to investigate antifungal tolerance in *C. auris* to all three classes of antifungal drugs used to treat invasive candidiasis. We find that 1) *C. auris* is tolerant to several common fungistatic and fungicidal drugs, which in some cases can be visually detected after 24 hours, as well as after 48 hours, of antifungal drug exposure; 2) the tolerant phenotype reverts to the susceptible phenotype in *C. auris*; and 3) combining azole, polyene, and echinocandin antifungal drugs with the adjuvant chloroquine reduces or eliminates tolerance and resistance in patient-derived *C. auris* isolates. These results suggest that tolerance contributes to treatment failure in *C. auris* infections for a broad range of antifungal drugs and that antifungal adjuvants may improve treatment outcomes for patients infected with antifungal-tolerant or antifungal-resistant fungal pathogens.

## 1. Introduction

Antimicrobial resistance (AMR) threatens the advances of modern medicine. Antifungal resistance contributes significantly to the AMR problem [1,2], especially among immunocompromised patients [3,4]. A multitude of biological, sociological, and economic factors result in hundreds of millions of serious fungal infections and between 1 and 1.5 million fungal infection-related deaths per year globally [5,6]. AMR among fungi is of particular concern due to the limited number of classes of drugs available to treat invasive fungal infections (i.e., fungistatic azoles, and fungicidal polyenes and echinocandins) [7]. This threat is exacerbated by the fact that no new class of antifungal drugs has reached the market in over a decade [8,9]. Climate change is also predicted to increase the prevalence of fungal infections, as fungi adapt to warmer temperatures to increase their geographic range and overcome the thermal protection barrier of their warm-blooded hosts [10].

*Candida* species of yeast are the most common causes of fungal infections [11]. One *Candida* species that is increasingly of concern is *Candida auris* [12] due to its resistance to antifungal drugs and healthcare-associated outbreaks [13]. *C. auris* has now been reported on all inhabited continents and in over 47 countries [14,15]. Particularly concerning is that *C. auris* is multidrug-resistant (i.e., non-susceptible to at least one agent in three or more classes of antimicrobials) [16–18], and, in some cases, it has been found to be pandrug-resistant (i.e., non-susceptible to all agents in all antimicrobial classes) [18,19]. *C. auris* has mortality rates of up to 45 % among patients with bloodstream infections [20].

“Tolerance” is a phenomenon whereby a slow growing subpopulation of cells, which are thought to be genetically identical to susceptible cells, emerges during antifungal drug treatment [21]. Antifungal tolerance is distinct from antifungal resistance, in that resistance is the result of heritable genetic changes and resistant cells grow above the minimum inhibitory concentration (MIC) in a concentration-dependent manner (i.e., MIC increases in resistance, but it does not increase in tolerance). In contrast, tolerance is a reversible phenomenon whereby cells grow slowly above MIC (i.e., they exhibit growth at “supra - MIC”). Tolerance manifests from the phenotypic heterogeneity intrinsic to a given fungal isolate, such that any cell within an isogenic population can reproduce the fractions of susceptible and tolerant cells present prior to the initiation of antifungal treatment. Cross tolerance has been observed in *C. albicans*, whereby strains tolerant to posaconazole also exhibit tolerance to other azole drugs [22]. Though the molecular mechanisms underlying tolerance in *Candida* species are still largely unknown, preliminary studies have shown that tolerance is associated with multiple genetic components that differ between isolates, including Hsp90-faciliated azole tolerance in *C. auris* [23]. Aneuploidy has also been shown to alter antifungal tolerance in *C. albicans* [24,25]. It is unknown if *C. auris* is tolerant to non-azole classes of antifungal drugs.

Clinical assays have not been designed to detect antifungal tolerance [26,27]. Quantitatively measuring tolerance of infecting isolates may provide prognostic insights concerning the success of mono- and combination-antifungal therapies [28]. Broth microdilution assays and disk diffusion assays coupled with the image analysis software *diskimageR* have been successfully used to quantify antifungal tolerance in research laboratories [29]. Most clinical diagnostic tests are performed on cultures grown for 24 hours and therefore cannot detect drug-tolerant cells, which are typically visually evident after 48 hours of growth [21]. Tolerance, along with host factors, immune status, and pharmacological issues [30], may explain why some patients do not respond to drug therapy despite being infected with fungi that have been determined by traditional antimicrobial susceptibility testing methods to be susceptible to a particular drug (i.e., cells that do not grow above MIC at 24 hours, the standard endpoint MIC measurement for *Candida* species) [21,28]. “Tailing growth” (the clinical term for tolerance) leads to poor response to fluconazole in *C. tropicalis* in wax moth larvae [31] and mouse models [32], and high levels of tolerance are associated with *C. albicans* infections in patients treated with fluconazole [33].

Adjuvant drugs have the potential to sustain the vital functions of antimicrobial drugs [34]. Non-antifungal agents have been shown to enhance the effectiveness of azole drugs against resistant *Candida* species and other pathogenic fungi, including *Aspergillus fumigatus, Cryptococcus neoformans*, and the dimorphic fungus *Histoplasma capsulatum* [35–37]. Specifically, the antimalarial drug chloroquine in combination with fluconazole exhibited enhanced antifungal activity against *C. albicans*, *C. tropicalis*, *C. glabrata, C. parapsilosis*, and *C. krusei (*teleomorph is known as *Issatchenkia orientalis* and *Pichia kudriavzevii* [11]) isolates *in vitro* [38]. If tolerance and resistance to azoles or to other classes of antifungal drugs can be eliminated in *C. auris* using adjuvant-antifungal therapies remains to be investigated. Another study explored the activity of doxycycline, pyrvinium pamoate, along with chloroquine, as adjuvants in combination with fluconazole in clinical *C. albicans* isolates and found increased antifungal activity [29]. Chloroquine is a member of quinoline family and is used to treat diseases including malaria, amebiasis, rheumatoid arthritis, discoid, and systemic lupus erythematosus [39–41]. Chloroquine causes iron depletion leading to a decrease in membrane sterol availability and downregulates the *ERG11* gene [42. We hypothesize that the combining chloroquine with common antifungal drugs will eliminate antifungal tolerance in *C. auris*.

The main aims of our study are to use broth microdilution and disk diffusion assays together with *diskImageR* to investigate if tolerance to all three classes of antifungal drugs occurs in *C. auris* and if this tolerance can be eliminated by adjuvant-antifungal therapy. We find that *C. auris* is tolerant to several fungistatic and fungicidal drugs: fluconazole, itraconazole, posaconazole, voriconazole, amphotericin B, and caspofungin. We demonstrate that antifungal tolerance is detectable at 24 hours, as well as at 48 hours, and that tolerance is a reversible phenomenon. Finally, we discover that combining antifungal drugs with the adjuvant chloroquine eliminates tolerance and resistance in *C. auris*.

## 2. Materials and Methods

### 2.1. Strains, Media, and Growth Conditions

*C. auris* isolates were obtained from clinical samples from the Alberta Precision Laboratories (APL) - Public Health Laboratory (ProvLab).

All strains and isolates (**Table A1**) were preserved in 25% glycerol at −80 °C until further use. The strains and isolates were revived by culturing from frozen stock on YPD agar plates (yeast extract: Sigma Aldrich, #8013-01-2; bacto peptone: Difco, #9295043) and incubated at 35 °C for 48 hours. Fresh subcultures were made on YPD agar plates and incubated at 35 °C for 24 hours prior to conducting microbroth dilution and disk diffusion assays (Section 2.3).

### 2.2. DNA Extractions, PCR, and Sequencing

The initial identification of all *C. auris* isolates was performed using matrix assisted laser desorption ionization – time of flight (MALDI-TOF) mass spectrometry [43,44] by the APL - ProvLab. The molecular identity of these isolates was confirmed by amplifying and sequencing the Internal Transcribed Spacer (ITS) region of ribosomal DNA. The primers ITS-5 (5’-GGAAGTAAAAGTCGTAACAAGG-3’) and ITS-4 (5’-TCCTCCGCTTATTGATATGC-3’) were used to amplify the ITS region (Integrated DNA Technologies). Genomic DNA was extracted using manual phenol-chloroform-isoamyl alcohol method [45]. The concentration of the extracted DNA was measured using a microvolume μDrop Plate (Thermo Fisher Scientific, #N12391). The template and the primers were mixed in concentration of 7.5 ng/μl and 0.25 μM, respectively, to a final volume of 10 μl. Sanger sequencing was then performed using a 3730 Genetic Analyzer (Thermo Fisher Scientific, #A41046) at the Molecular Biology Services Unit at the University of Alberta. The resulting sequences were subjected to nucleotide BLAST analysis [46], which revealed 100 % similarly to the standard strains. The *C. auris* isolates’ ITS sequences were submitted to NCBI with the accession number OP984814-OP984818.

### 2.3. Broth Microdilution and Disk Diffusion Assays

The MIC for each isolate was first determined via broth microdilution assays following CLSI M27 guidelines [47]. All the isolates were tested in 96-well U-bottom microwell plates (Thermo Fisher Scientific, #163320) against fluconazole (Sigma-Aldrich, #F8929) (0.12-64 μg/ml), amphotericin B (Sigma-Aldrich, #A9528) (0.03-16 μg/ml), itraconazole (Sigma-Aldrich, #16657) (0.03-16 μg/ml), posaconazole (Sigma-Aldrich, #SML2287) (0.03-16 μg/ml), voriconazole (Sigma-Aldrich, #P20005) (0.03-16 μg/ml), micafungin (Sigma-Aldrich, #208538) (0.015-8 μg/ml), caspofungin (Sigma-Aldrich, #179463-17-3) (0.015-8 μg/ml), and anidulafungin (Sigma-Aldrich, #166663-25-8) (0.03-16 μg/ml). These antifungals were dissolved in DMSO (fluconazole, itraconazole, voriconazole, posaconazole, and amphotericin B) or water (caspofungin and micafungin); the concentration of the antifungal microwell plates were twice the final concentration tested with the inoculum added. Freshly cultured *Candida* species (*C. auris*, *C. parapsilosis* (ATCC 22019), and *I. orientalis* (ATCC 6258)) at 24 hours of incubation at 35 °C were used as inoculum. Inoculum of 100 μl consisting of 2-5 x 10^3^ cells were used to inoculate the antifungal microwell plates. After inoculation the microwell plates were incubated at 35 °C and evaluated after 24 hours and 48 hours to determine the MICs.

Disk diffusion assays were carried out as per CLSI M44-A2 guidelines [48] against fluconazole (25 μg), itraconazole (50 μg), posaconazole (5 μg), voriconazole (1 μg), amphotericin B (20 μg), and caspofungin (5 μg). MHA medium with 2 % dextrose (Sigma Aldrich, #50-99-7) and 0.5 μg/mL methylene blue dye (Sigma Aldrich, #03978) was used to perform the disk diffusion assays. After 24 hours of growth, 5-10 colonies were picked and liquid suspensions of *C. auris* were made by reconstituting colonies in 2 ml of normal saline (Sigma Aldrich, #S8776). The optical density (OD) was measured using a Varioskan LUX microplate reader (Thermo Fisher Scientific, #N16044) at 530 nm and adjusted to an OD of 0.09-0.13, which corresponded to 1-5 x 10^6^ cells/mL. The adjusted solution was utilized to swab on the Muller-Hinton agar (MHA) using sterile cotton swabs (Fisher Scientific, #22-029-683). An antifungal disk was placed on each plate after inoculating and drying the agar plates. The plates were then incubated for 24 to 48 hours at 35 °C. All experiments were performed in triplicate.

### 2.4. Photography and Image Preprocessing

Photographs of each disk diffusion plate were taken after 24 hours and 48 hours at the maximum possible resolution (6000 by 4000 pixels with an aspect ratio of 3:2) using a Canon EOS Rebel SL3 camera with a Canon EF-S 35mm f/2.8 Macro IS STM macro lens. The camera settings were as follows: ISO 800, white balance, picture type “neutral”, time 1/100 s, center focused against a plain black background from a fixed distance. The photos were taken and then the size of each photograph was standardized by cropping the edges and bringing all images to the same resolution.

### 2.5. Quantifying Tolerance via Supra-MIC Growth and Fraction of Growth

Tolerant subpopulations grow slowly in drug concentrations above MIC [21]. We used established methods to quantify tolerance, namely, supra-MIC growth from microbroth dilution assays and the fraction of growth (FoG) in the zone of inhibition (ZOI) from disk diffusion assays (Section 2.3).

The MIC for each isolate was determined using CLSI supplement M60 guidelines [49]. The MIC readings were recorded at 24 hours and 48 hours post inoculation. Tentative breakpoints provided by the Centers for Disease Control and Prevention for *C. auris* were considered to differentiate them as susceptible or resistant [50]. *I. orientalis* and *C. parapsilosis* were used as reference strains to ensure that the antifungal MIC range in each experiment was within CLSI guidelines.

Supra-MIC growth (SMG) was determined by subjecting the antifungal microwell plates used for measuring MICs by spectrophotometric reading at 630 nm after 24 hours and 48 hours of incubation at 35 °C. SMG was calculated as an average growth per well above MIC normalized to total growth without antifungals [28]:

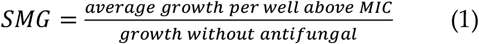

The software program *diskImageR* [29] analyses photographs of disk diffusions assays. *diskImageR* utilizes the image processing program *ImageJ* [51] and the programming language *R* [52]. We used *diskImageR* to measure the tolerance and resistance of *C. auris* isolates to fungistatic and fungicidal drugs from photographs of the disk diffusion assay plates (Section 2.4; **Figures B1 and B2**). All disk diffusion experiments were repeated in triplicate using antifungal disks placed in the center of MHA plates incubated at 35 °C for 24 and 48 hours (**Figure B3**). After the photographs were imported by *diskImageR* into *ImageJ*, the size of each photograph was standardized and the ‘find particles’ macro was used to find the center of the antimicrobial diffusion disk. The radius of the ZOI (RAD) and the FoG in the ZOI were measured where 20 %, 50 %, and 80 % of growth was inhibited (RAD_20_, RAD_50_, and RAD_80_, and FoG_20_, FoG_50_, and FoG_80_, respectively). The RAD measures the degree of susceptibility/resistance and FoG measures the degree of tolerance. The RAD for all disk diffusions assay plates were also measured manually (using a ruler) and the FoGs were also analyzed using *ImageJ* [53]. *ImageJ* analysis for estimating pixel intensity to obtain FoG was carried out by importing photographs to *ImageJ* software and setting ‘on’ the measurements such as ‘mean grey value’, ‘minimum and maximum grey ‘area’, and fixing the ‘area’ for ZOI. ‘Measure’ macro was then used to measure the pixel intensity. For photographs of 48 hours DDA plates, same parameters were restored as for its 24 hours counterpart and the pixel intensity was measured within ZOI. When there are colonies at border of the ZOI (e.g., **Figure B3B**), *diskImageR* considers it as the area outside of the ZOI, and the measured RAD is smaller than the manually measured RAD; consequently, the FoG measured by *diskImageR* is also inaccurate. Therefore, in these cases, the RAD was obtained by manually measuring the RAD and by measuring the FoG using *ImageJ* (**Figure B4**) [51]. When isolates were highly tolerant, resulting in many colonies in the ZOI (**Figure B5B**) or complete confluence in the ZOI (**Figure B5D**) after 48 hours, *diskImageR* reported RAD and FoG as “NA” (Not Applicable).

### 2.6. Experiments to Determine Effectiveness Adjuvant-Antifungal Treatment

The synergy among antifungals (fluconazole, itraconazole, posaconazole, voriconazole, amphotericin B, and caspofungin) and adjuvant (chloroquine) against *C. auris*, *C. parapsilosis*, and *I. orientalis* were evaluated using DDAs (Section 2.3) with minor modifications. Syringe filtered chloroquine diphosphate salt (Sigma-Aldrich, #C6628) solution was added to MHA media after autoclaving to a final concentration of 1,031.8 μg/mL. After inoculation of *C. auris* and the control strains, the MHA plates containing chloroquine were incubated in dark as chloroquine light sensitive. These plates were read and photographed at 24 hours and 48 hours. *C. auris* isolates and control strains were lawn cultured (i.e., the entire surface of the agar plate was covered by swabs dipped in the liquid culture) on the MHA plates containing chloroquine with and without antifungal disks to respectively determine the effect of antifungal-chloroquine and chloroquine alone on *C. auris*.

## 3. Results

### 3.1. Identification of Resistance in C. auris from Broth Microdilution Assays

To determine if the *C. auris* isolates were resistant to the antifungal drugs used in our study, we performed antifungal susceptibility testing at 24 and 48 hours using the broth microdilution method (Section 2.3). The MICs for the *C. auris* isolates indicated that all isolates were susceptible to the fungicidal and fungistatic drug tested, except for *C. auris* isolate 2 which was not susceptible to fluconazole and *C. auris* isolate 5 which was not susceptible to fluconazole, voriconazole, caspofungin, and amphotericin B (**Figure 1** and **Table A2**). The quality control strains *C. parapsilosis* and *I. orientalis* were within the recommended range. No change in MIC was witnessed at 24 and 48 hours except for *C. auris* isolate 1 and 2 against amphotericin B.

**Figure 1.**
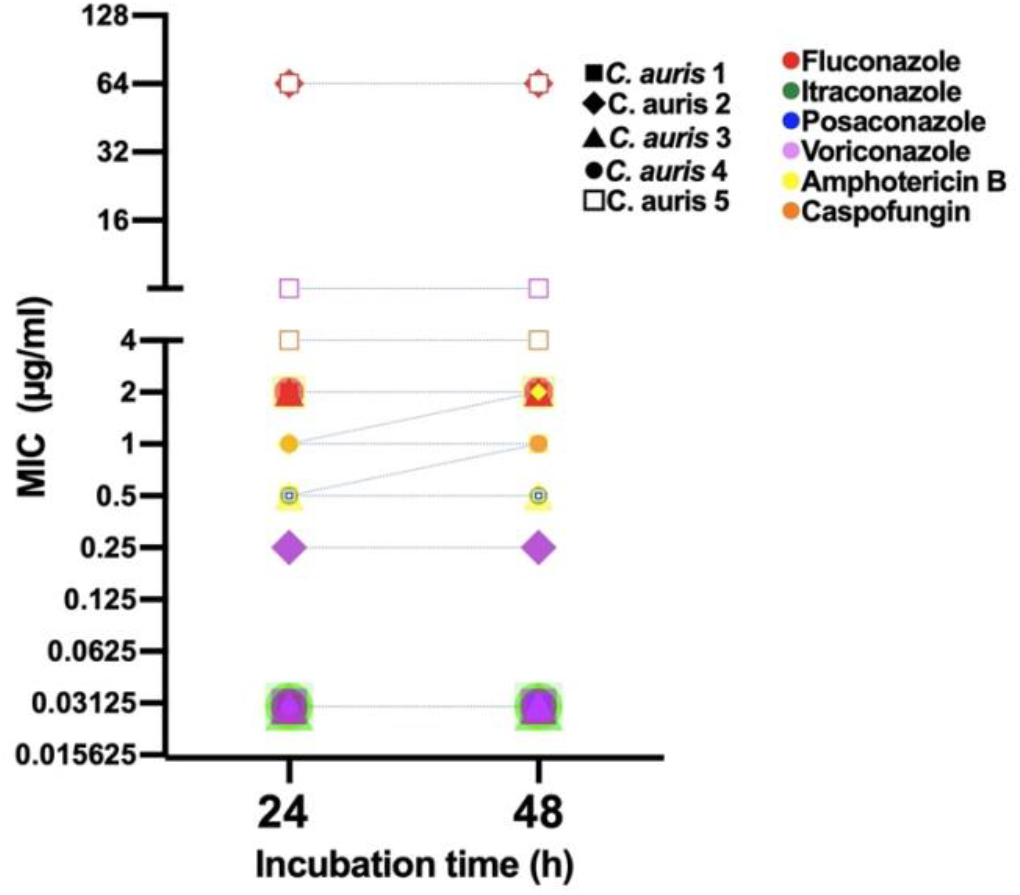
Minimum inhibitory concentration (MIC) for clinical *C. auris* isolates growing in antifungal microwell plates to determine susceptibility/resistance to antifungal drugs. Mean MICs of five clinical *C. auris* isolates measured after 24 and 48 hours for four fungistatic drugs (fluconazole, itraconazole, posaconazole, and voriconazole) and two fungicidal drugs (amphotericin B and caspofungin).

### 3.2. Identification of Resistance in C. auris from Disk Diffusion Assays

To confirm the resistance of the *C. auris* isolates determined by the broth microdilution assays (Section 3.1), we performed the corresponding disk diffusion assays. In agreement with the microbroth dilution method, resistance was noted in *C. auris* isolate 2 for fluconazole and *C. auris* isolate 5 for fluconazole and voriconazole (RAD = 0 mm in all three instances; **Figure 2A**). However, *C. auris* isolate 5 exhibited a ZOI to amphotericin B (RAD = 10 mm) and caspofungin (RAD = 6 mm) (**Figure 2A**). As expected, and in agreement with previous work [28], there was an inverse correlation between RAD and MIC (Pearson test, *r* = −0.58, *p* = 0.007).

**Figure 2.**
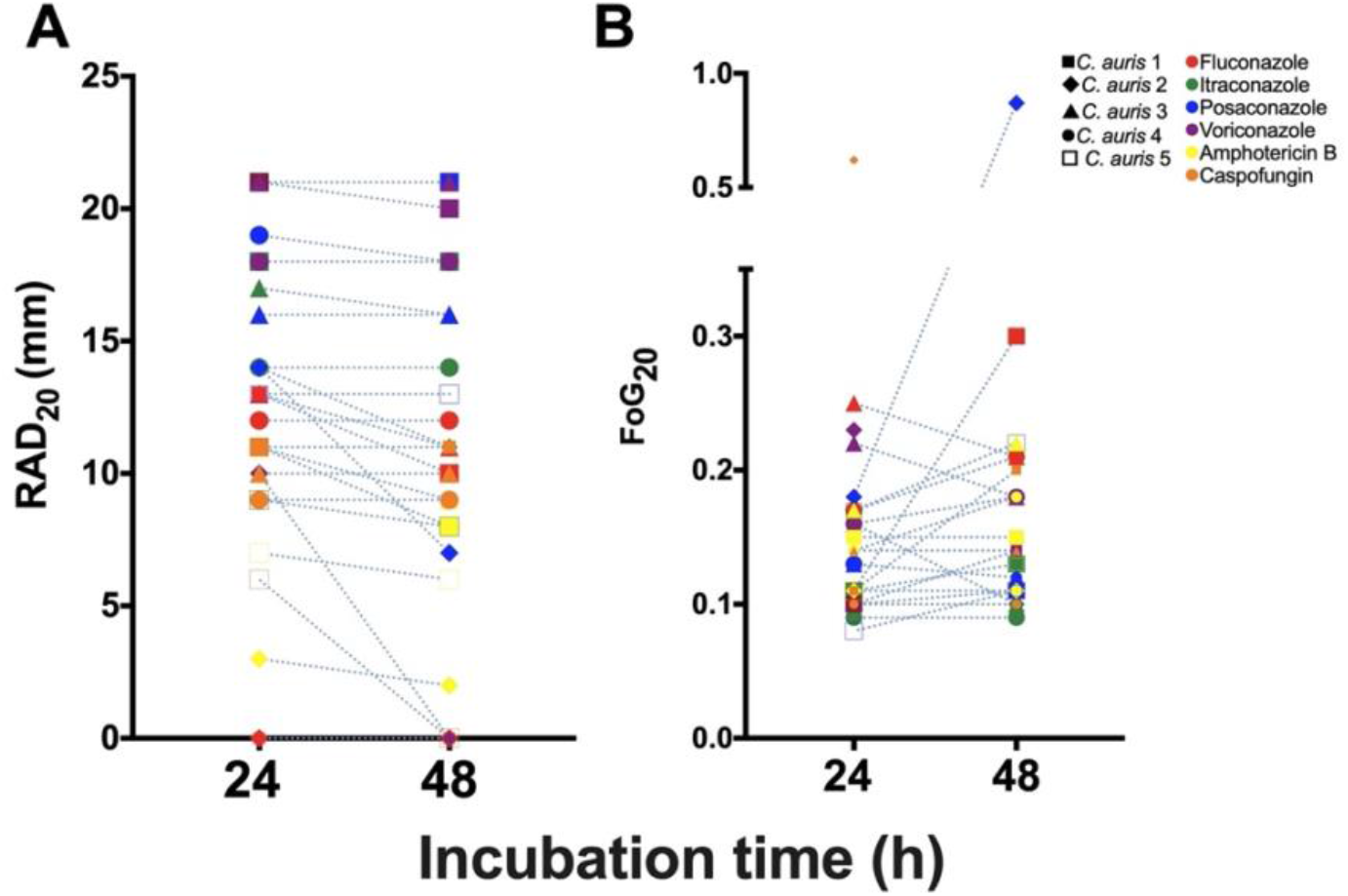
Radius of the zone of inhibition (RAD) and fraction of growth in the zone of inhibition (FoG) for *C. auris* isolates treated with antifungal drugs. **(A)** Mean RAD where 20 % of growth is inhibited (RAD20) at 24 and 48 hours. **(B)** Mean FoG where 20 % of growth is inhibited (FoG20) 24 and 48 hours. *C. auris* isolate 2 treated with caspofungin at 48 hours is not plotted in (B) as it exhibited FoG in entire ZOI (i.e., a “NA” data point was generated by *diskImageR* [29]); the reduction in RAD and FoG for *C. auris* isolate 3 in (A) and (B), respectively, is due to the exclusion of FoG within the ZOI by *diskImageR* (see Section 2.5 for details).

### 3.3. Identification of Tolerance in C. auris from Broth Microdilution Assays

To determine if tolerant subpopulations existed within the non-resistant *C. auris* isolates, we carried out an SMG analysis (Section 2.5). A statistically significant increase in SMG was observed after 48 hours for *C. auris* isolate 1 to fluconazole and itraconazole (Independent t-test, *p* = 0.009 and *p* = 0.001, respectively), *C. auris* isolates 2 and 4 to voriconazole (Independent t-test, *p* = 0.0009 and *p* = 0.0014, respectively), and *C. auris* isolate 2 to caspofungin (Independent t-test, *p* = 0.006) indicating the presence of tolerance (**Figure 3A**). There was also a non-significant increase in SMG at 48 hours for *C. auris* isolates 3 and 4 to fluconazole, *C. auris* isolates 2, 3, and 5 to itraconazole, *C. auris* isolates 1, 2, 3, 4, and 5 to posaconazole, *C. auris* isolates 1 and 3 to voriconazole, and *C. auris* isolate 1 to amphotericin B. No tolerance was observed for *C. auris* isolate 4 to itraconazole and caspofungin, and *C. auris* isolates 2, 3, and 4 to amphotericin B (**Figure 3B**). There was a decrease in SMG for *C. auris* isolate 1 against amphotericin B. This occurred because the growth of isolates in wells without antifungals increased over 48 hours, which in turn reduced the SMG (as described in **Equation 1**).

**Figure 3.**
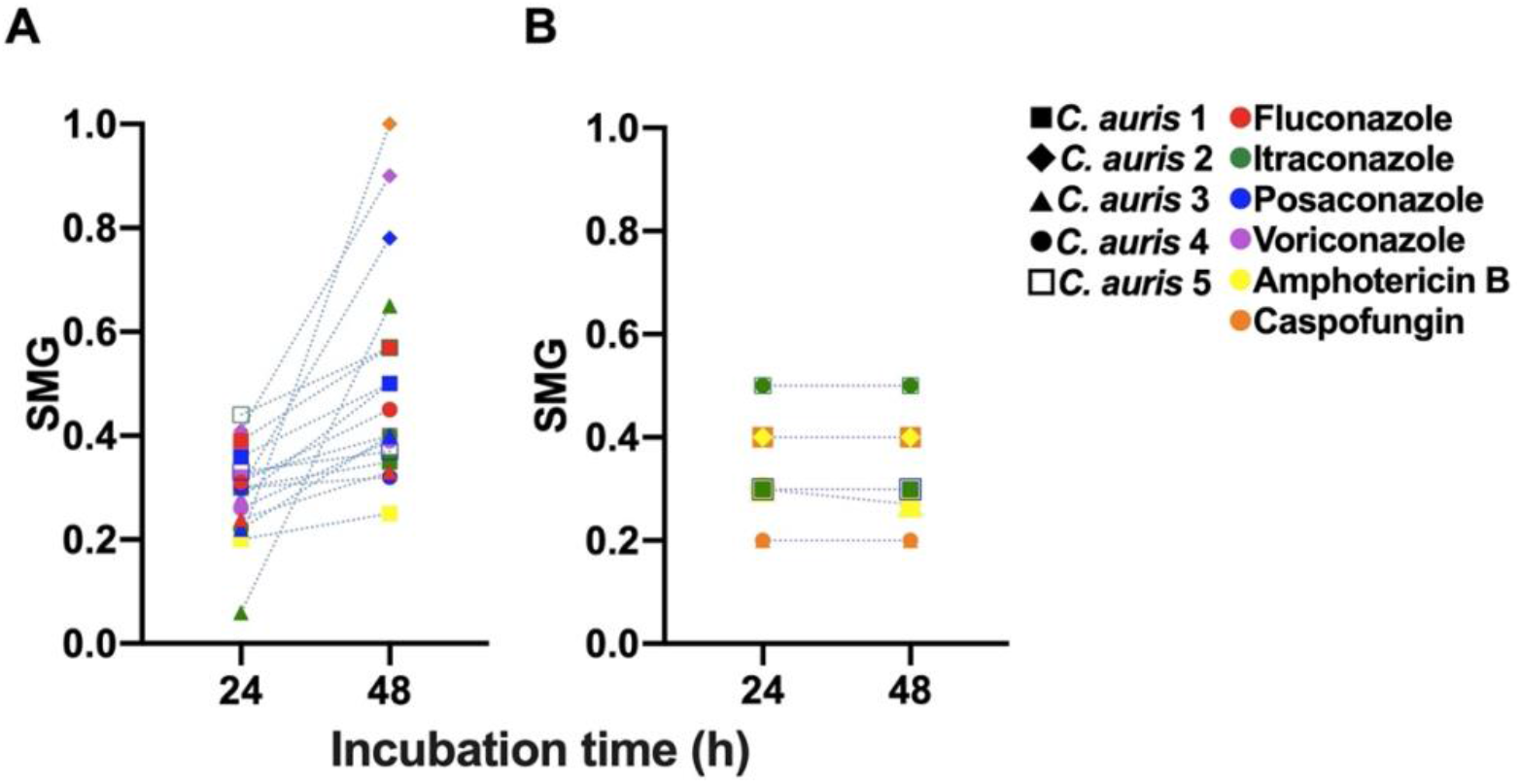
Tolerance from supra-MIC growth (SMG) for clinical *C. auris* isolates grown in antifungal microwell plates. **(A)** Mean SMG of tolerant isolates after 24 and 48 hours. **(B)** Mean SMG of non-tolerant isolates after 24 and 48 hours.

### 3.4. Identification of Tolerance in C. auris from Disk Diffusion Assays

To confirm the tolerance of the *C. auris* isolates determined by the broth microdilution assays (Section 3.3), we performed the corresponding DDAs. All the *C. auris* isolates with higher SMG exhibited higher FoG at 48 hours. The FoG within the ZOI ranged between 0.08 - 0.62 and 0.09 - 0.87 at 24 hours and 48 hours, respectively (**Figure 2B**). *C. auris* isolate 2 exhibited the highest FoG against caspofungin at 24 hours (0.62) and against posaconazole at 48 hours (0.87). Similarly, at 24 hours the highest pixel intensity occurred for *C. auris* isolate 3 against posaconazole (195, **Figure 4A**) and the highest SMG occurred for *C. auris* isolate 2 against caspofungin (1.0, **Figure 4A**). At 48 hours the highest pixel intensity and SMG were measured for *C. auris* isolate 2 against voriconazole (197 and 0.90, respectively; **Figure 4B**).

**Figure 4.**
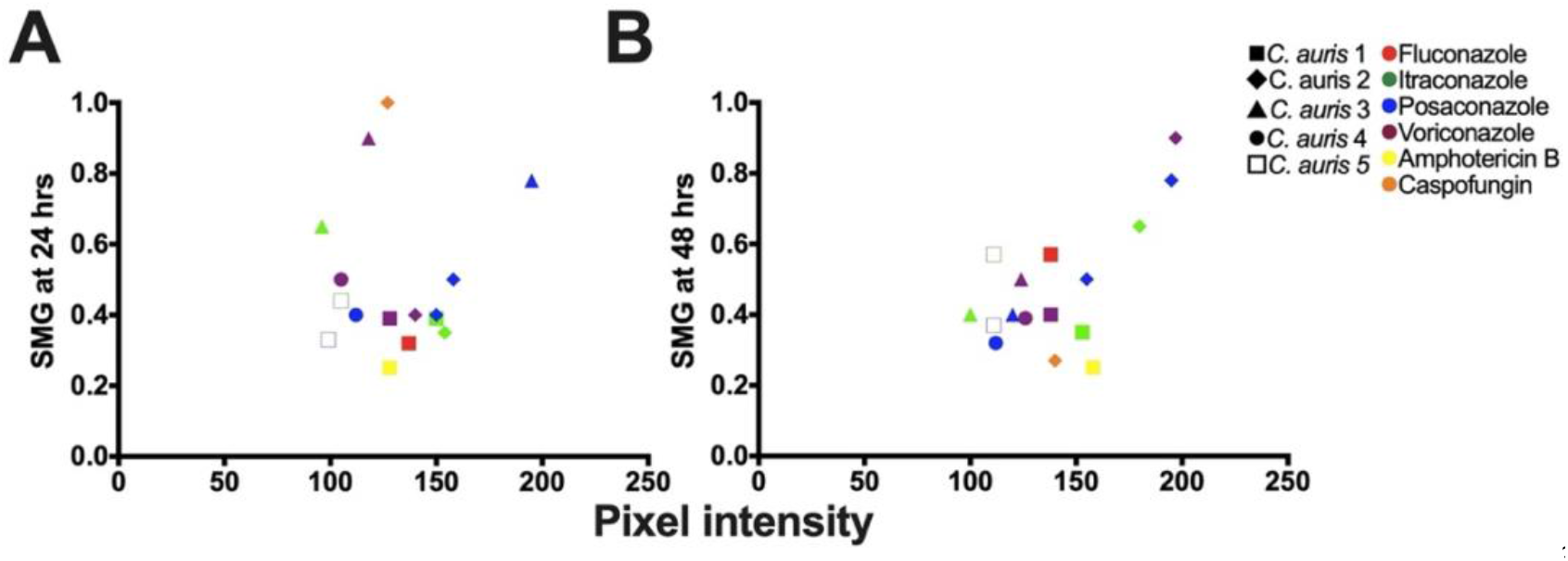
Correlation analysis for mean supra-MIC growth (SMG) and mean pixel intensity measured by *ImageJ* [51] to determine tolerance. **(A)** Analysis performed after 24 hours of growth (*R^2^* = 0.3128; Pearson correlation test, *p* = 0.0469). **(B)** Analysis performed after 48 hours of growth (*R^2^* = 0.2862; Pearson correlation test, *p* = 0.0085).

There was no correlation between FoG and RAD levels (Pearson test, *r* = −0.25, *p* = 0.28), as expected based on previous work that established the FoG and RAD measure different drug responses [28,29]. The was significant correlation between SMG measured by *diskImageR* and pixel intensity measured by *ImageJ* (**Figure 4**), which occurred as both SMG and pixel intensity increase when tolerant subpopulations are present.

Overall, there was no significant difference between *diskImageR* and manual readings of the RAD (Independent t-test, *p* = 0.5634 and *p* = 0.8453 for readings at 24 hours and 48 hours, respectively; **Figure B4**). There was also no significant difference for FoG readings using *diskImageR* and *ImageJ* at 24 hours (Unpaired t-test, *p* = 0.35). However, there was a statistically significant difference for FoG reading using *diskImageR* and *ImageJ* at 48 hours (Unpaired t-test, *p* = 0.022). The difference these FoG readings resulted from the fact that *diskImageR* was unable to distinguish the border of the ZOI among tolerant isolates, which was obscured by tolerant colonies at 48 hours.

Among reference strains, only *C. parapsilosis* exhibited tolerance to fluconazole and voriconazole (**Figure B6**). The FoG and SMG for fluconazole and voriconazole is presented in **Table A3**. No tolerance was observed for the other antifungal drugs considered in this study against *C. parapsilosis*. *I. orientalis* did not exhibit tolerance to any of the antifungal agents tested.

### 3.5. Tolerance in C. auris is a Reversable Phenomenon

Next, we investigated if the antifungal tolerance that we discovered in *C. auris* was a reversible phenomenon. To investigate this, we sub-cultured colonies growing within and outside of the ZOI and repeated the microbroth dilution and disk diffusion experiments (**Figure B7**). There was no difference between the MICs of original colonies and colonies from inside and outside ZOI at both 24 and 48 hours (**Table A4**). RAD, FoG, and SMG, obtained from *C. auris* colonies isolated from inside and outside the ZOI also did not show any statistically significant differences. These results indicate that the antifungal-tolerant colonies our experiments could reversibly generate antifungal-susceptible colonies.

### 3.6. Elimination of Tolerance and Resistance in C. auris via Adjuvant-Antifungal Treatment

To eliminate the tolerance observed in our clinical *C. auris* isolates (Sections 3.3 and 3.4) a previously known adjuvant chloroquine [38] was combined with the antifungal drugs fluconazole, itraconazole, posaconazole, voriconazole, amphotericin B, and caspofungin. Chloroquine-antifungal disk diffusion assays were performed on all five clinical *C. auris* isolates, as well as on the *C. parapsilosis* and *I. orientalis* reference strains (**Table A1**). Chloroquine alone did not have any antifungal effect on either *C. auris* isolates or the reference strains (**Figure B9**).

Tolerance and resistance were reduced or eliminated in our clinical *C. auris* isolates by combing chloroquine with antifungal drugs. *C. auris* isolate 1 showed increase in RAD for fluconazole, posaconazole, amphotericin B, and caspofungin in presence of chloroquine compared to RAD measured with these antifungal drugs alone (**Figure 5A,B**). Similarly, *C. auris* isolate 2 for posaconazole, amphotericin B, and caspofungin; *C. auris* 3 for posaconazole, voriconazole, amphotericin B, and caspofungin; *C. auris* isolate 4 for itraconazaole, posaconazole, voriconazole, amphotericin B, and caspofungin; and *C. auris* isolate 5 for itraconazole and caspofungin (elimination of resistance for caspofungin) all displayed a reduction in RAD when these antifungal drugs were combined with chloroquine. Correspondingly, the FoG was reduced in presence of chloroquine for *C. auris* isolate 1 when combined with posaconazole, amphotericin B, and caspofungin (**Figure 5C,D**). Similar adjuvant-antifungal FoG results were obtained for *C. auris* isolate 2 against posaconazole, voriconazole, amphotericin B and caspofungin; *C. auris* isolate 3 against fluconazole, itraconazole, posaconazole, voriconazole, amphotericin B, and caspofungin; *C. auris* isolate 4 against amphotericin B; and *C. auris* isolate 5 against itraconazole and caspofungin. The FoG for *C. auris* isolate 2 for voriconazole and caspofungin and *C. auris* isolate 5 for caspofungin could not be measured at 48 hours without chloroquine as there was no ZOI. However, it was possible in presence of chloroquine indication the presence of ZOI and reduction in the tolerance as well as resistance. The reference strain *I. orientalis* (resistant to fluconazole) exhibited a ZOI against fluconazole when supplemented with chloroquine (**Table A5; Figure B8**), however, *C. parapsilosis* was not significantly affected by the presence of chloroquine **(Table A5)**.

**Figure 5.**
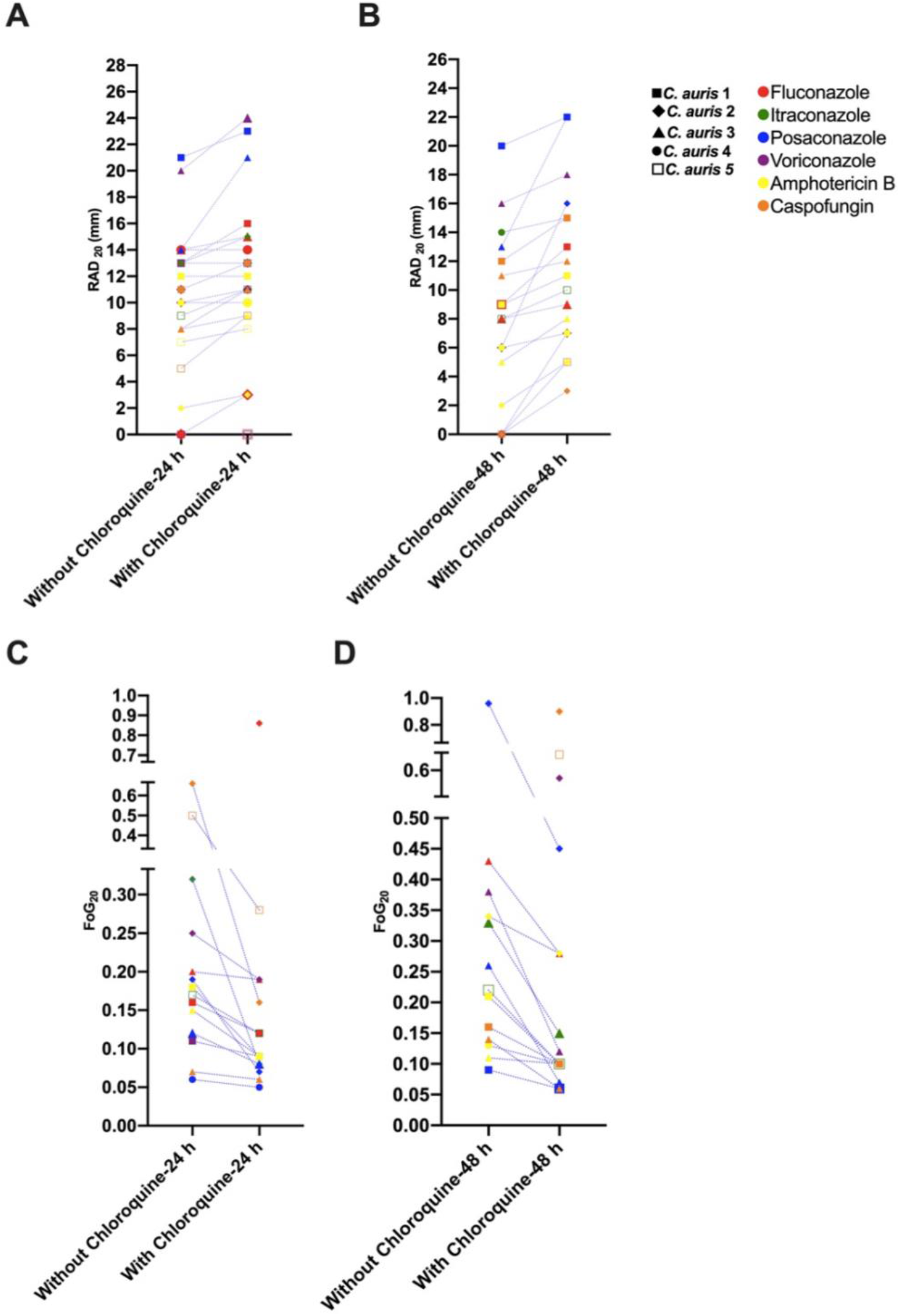
Radius of the zone of inhibition (RAD) and fraction of growth in the zone of inhibition (FoG) measurements for *C. auris* isolates for adjuvant -antifungal disk diffusion assays. **(A)** Mean RAD measured for the *C. auris* isolates at 24 hours against antifungal drugs with and without the adjuvant chloroquine. **(B)** Mean RAD measured for the *C. auris* isolate at 48 hours against antifungal drugs with and without chloroquine. **(C)** Mean FoG measured using *diskImageR* [29] for all *C. auris* isolates at 24 hours against antifungal drugs with and without chloroquine. **(D)** Mean FoG measured using *diskImageR* for the *C. auris* isolates at 48 hours against antifungal drugs with and without chloroquine. Note that the single data points in (C) and (D) at 48 hours are due to the mitigation of resistance in presence of chloroquine, as FoG could not be measured for these isolates at 24 hours because of their resistance to the corresponding antifungal drugs.

## 4. Discussion

We discovered that *C. auris* is tolerant to fungistatic drugs (fluconazole, voriconazole, itraconazole, and posaconazole) and to fungicidal drugs (amphotericin B and caspofungin). We also found azole tolerance in *C. parapsilosis* (fluconazole and voriconazole) but not in *I. orientalis*; *I. orientalis* was intrinsically resistant to fluconazole. We were able to visually detect tolerance after 24 hours, as well as after 48 hours by FoG, of antifungal treatment using *diskImageR* [29] and *ImageJ* [53]. These finding suggests that a distinct subpopulation among *C. auris* were able to survive and grow slowly in the presence of different antifungal drugs. Since, *C. auris* is a multidrug-resistant pathogen, the presence of tolerance further narrows treatment options. Previous reports suggest that tolerant subpopulations among infecting *Candida* species are strongly associated with mortality among candidemia patients [54]. Therefore, clinical diagnostic laboratories should also test for antifungal tolerance along with standard antifungal susceptibility/resistance tests to increase the efficacy of antifungal treatment. Furthermore, existing tolerance quantification methods could be adapted to detect tolerance after 24 hours and 48 hours to broaden the scope of standard antimicrobial susceptibility testing in medical diagnostic laboratories. The fluconazole tolerance that we observed in *C. auris* was in agreement with previous studies on *C. albicans* [28] and *C. auris* [23], as well as with related clinical studies on “trailing growth” (reduced but persistent visible growth of *Candida* species in fluconazole concentrations above MIC [32,55,56]).

The tolerance to fungistatic and fungicidal drugs observed in the clinical *C. auris* isolates our study appears to be a reversible phenomenon, as previously described for clinical *C. albicans* isolates [57]. The tolerant cells growing inside ZOI upon sub-culture are indistinguishable from parental population suggesting the presence of phenotypic heterogeneity instead of genetic variations. *C. auris* isolates cultured from inside and outside the ZOI did not show any significant changes in the RAD, MIC, or SMG levels. This reversible tolerance that we observed in *C. auris* may result from stochastic phenotype switching or an induced response activated by the presence of antifungal drugs inside of the cell. The general mechanism underlying tolerance in *C. auris* remains to be elucidated in future work, to be aided for instance, by mathematical modeling and synthetic biology [58], tracking single cell growth and gene expression trajectories in microfluidic devices [59,60], as well as genetic sequence and aneuploidy analyses [61].

The tolerance in our *C. auris* isolates was reduced or eliminated *in vitro* by combining azole, polyene, and echinocandin antifungal drugs with the antimalarial drug chloroquine. Combining chloroquine with antifungal drugs had a similar effect on resistance on the *C. auris* isolates investigated in this study. *C. auris* isolate 5, which was resistant to caspofungin and voriconazole (RAD = 0 mm), had a small increase in the ZOI (RAD < 12 mm) in presence of chloroquine. Correspondingly, *C. auris* isolate 2 had no ZOI for caspofungin, but had a small ZOI (RAD = 6) in presence of chloroquine. The RADs for these cases were smaller than those for the sensitive *C. auris* isolates in our experiments. However, a previous study classified *C. auris* as resistant to fluconazole and caspofungin when the ZOI was less than 12 mm, as this corroborated with their MIC results for resistant isolates [62]. Similarly, chloroquine also effects fluconazole resistance in *I. orientalis* (**Table A5**), though tolerant subpopulations in *C. parapsilosis* were unaffected by chloroquine. Altogether, these results suggest that combining chloroquine with antifungal drugs has a partial mitigation effect on resistance in *C. auris*. Though the mechanism of action is unknown, it is likely related to iron depletion caused by chloroquine cause and/or downregulation of the *ERG11* gene [36,42]. Iron depletion is known to decrease membrane sterols and increase membrane fluidity, leading to increased uptake of antifungal drugs into the cell [63]. The downregulation of *ERG11* gene, which synthesis lanosterol alpha demethylase enzyme, is also known to be an important rate-limiting enzyme for the synthesis of ergosterol [64].

Overall, this study advances our understanding of antifungal treatment failure in *C. auris* and identifies opportunities for the clinical detection of antifungal tolerance as well as the development of targeted adjuvant-antifungal therapies against tolerant and resistant invasive candidiasis.

## Supplementary Materials

This manuscript contains the following supporting materials: Table A1: *Candida* isolates and strains; Table A2: Minimum inhibitory concentrations (MICs) of *C. auris* isolates; Table A3: Mean MIC, SMG, FoG, and RAD for reference strains *Issatchenkia orientalis* and *C. parapsilosis* measured at 24 and 48 hours for different antifungal drugs. Table A4: Reversibility of tolerance phenotype in *Candida auris*. Table A5: Effect of chloroquine (CLQ) on reference strains *Issatchenkia orientalis* and *Candida parapsilosis*. Figure B1: Quantification of antifungal tolerance in a disk diffusion assay using the image analysis program *diskImageR* [29]. Figure B2: Detecting tolerance in *Candida auris* from disk diffusion assays (DDAs) using *diskImageR*. Figure B3: Representative disk diffusion assays (DDA) images of fluconazole (FLU) tolerance in *Candida auris* and *Candida parapsilosis*. Figure B4: Comparison between *diskImageR* and manual radius of the zone of inhibition (RAD) measurements. Figure B5: Azole tolerance in *Candida auris*. Figure B6: Azole tolerance in *Candida parapsilosis* reference strain. Figure B7: Reversibility of tolerance in a representative *Candida auris* isolate 2 against voriconazole. Figure B8: Disk diffusion assays (DDAs) of antifungal-adjuvant treatment in *Candida auris* isolates and *Issatchenkia orientalis* and *Candida parapsilosis* reference strains. Figure B9: *Candida auris* isolates and *Candida parapsilosis* and *Issatchenkia orientalis* reference strains growing on Mueller-Hinton agar (MHA) media with chloroquine.

## Author Contributions

Conceptualization, D.C.; methodology, S.S., S.R.K., and D.C.; experimental investigation, S.R.K., S.S., and C.M.G.; formal analysis and visualization, S.R.K. and S.S.; data curation, S.S. and S.R.K.; writing, original draft preparation, review, and editing, D.C., S.S, S.R.K., and C.M.G.; supervision, D.C. and S.S.; funding acquisition, D.C. All authors have read and agreed to the published version of the manuscript.

## Data Availability Statement

All data supporting the findings of this study are presented within the manuscript.

## Acknowledgments

We thank Dr. Tanis Dingle and Alberta Precision Laboratories - Public Health Laboratory for providing the *C. auris* isolates. We acknowledge the Molecular Biology Services Unit at the University of Alberta for assistance with genetic sequencing. Finally, we thank the late André T. Charlebois (1953-2020) for guidance on the camera setup and for advice on the macro photography of yeasts.

## Appendix A Supplementary Tables

**Table A1.** *Candida* isolates and strains. *Candida auris* and *Candida* reference strains *Issatchenkia orientalis* and *Candida parapsilosis* used in our study to investigate antifungal tolerance and resistance. *Issatchenkia orientalis* is also known by the binomial names *Candida krusei* and *Pichia kudriavzevii*. ID = identity number.

| Strain/Isolate Number | Genus | Species | ID |
| --- | --- | --- | --- |
| 1 | *Candida* | *auris* | RN190002998 |
| 2 | *Candida* | *auris* | RN180001039 |
| 3 | *Candida* | *auris* | 21UA-342M00567 |
| 4 | *Candida* | *auris* | 21DY-278M00215 |
| 5 | *Candida* | *auris* | 0579 |
| 6 | *Issatchenkia* | *orientalis* | ATCC 6258 |
| 7 | *Candida* | *parapsilosis* | ATCC 22019 |

**Table A2.** Minimum inhibitory concentrations (MICs) of *C. auris* isolates. Mean MICs were measured in μg/ml after 24 and 48 hours for a range of fungistatic and fungicidal drugs.

| Antifungal drug | Hours of incubation | *C. auris* 1 | *C. auris* 2 | *C. auris* 3 | *C. auris* 4 | *C. auris* 5 |
| --- | --- | --- | --- | --- | --- | --- |
| Fluconazole | 24 | 2 | 64 | 2 | 2 | 64 |
|  | 48 | 2 | 64 | 2 | 2 | 64 |
| Voriconazole | 24 | 0.03 | 0.25 | 0.03 | 0.03 | 8 |
|  | 48 | 0.03 | 0.25 | 0.03 | 0.03 | 8 |
| Itraconazole | 24 | 0.03 | 0.25 | 0.03 | 0.03 | 0.5 |
|  | 48 | 0.03 | 0.25 | 0.03 | 0.03 | 0.5 |
| Posaconazole | 24 | 0.03 | 0.25 | 0.03 | 0.03 | 0.5 |
|  | 48 | 0.03 | 0.25 | 0.03 | 0.03 | 0.5 |
| Amphotericin B | 24 | 0.5 | 1 | 0.5 | 1 | 2 |
|  | 48 | 1 | 2 | 0.5 | 1 | 2 |
| Caspofungin | 24 | 1 | 0.5 | 0.5 | 0.5 | 4 |
|  | 48 | 1 | 0.5 | 0.5 | 0.5 | 4 |
| Anidulafungin | 24 | 0.03 | 0.25 | 0.03 | 0.06 | 2 |
|  | 48 | 0.03 | 0.25 | 0.03 | 0.06 | 2 |
| Micafungin | 24 | 0.12 | 0.06 | 0.06 | 0.12 | 0.5 |
|  | 48 | 0.12 | 0.06 | 0.06 | 0.06 | 0.5 |

**Table A3.** Mean MIC, SMG, FoG, and RAD for reference strains *Issatchenkia orientalis* and *C. parapsilosis* measured at 24 and 48 hours for different antifungal drugs. MIC: minimum inhibitory concentration; SMG: Supra-MIC Growth; FoG: Fraction of growth; RAD: Radius of the zone of inhibition; NA: Not available.

|  |  | MIC<br>(µg/ml) | SMG<br>at 24 hours | SMG<br>at 48 hours | FoG<br>at 24<br>hours | FoG<br>at 48<br>hours | RAD<br>at 24<br>hours<br>(mm) | RAD<br>at 48<br>hours<br>(mm) |
| --- | --- | --- | --- | --- | --- | --- | --- | --- |
| *I. orientalis*<br>ATCC 6258 | Fluconazole | 32 | NA | NA | 0 | 0 | 0 | 0 |
|  | Itraconazole | 0.12 | 0.62 | 0.62 | 0.09 | 0.09 | 14 | 14 |
|  | Voriconazole | 0.25 | 0.65 | 0.25 | 0.09 | 0.1 | 12 | 12 |
|  | Posaconazole | 0.06 | 0.16 | 0.5 | 0.09 | 0.11 | 12 | 12 |
|  | Amphotericin B | 1 | 0.31 | 0.25 | 0.2 | 0.06 | 6 | 3 |
|  | Caspofungin | 0.5 | 0.62 | 0.5 | 0.13 | 0.22 | 11 | 11 |
| *C. parapsilosis*<br>ATCC 22019 | Fluconazole | 2 | 0.41 | 0.47 | 0.13 | 0.13 | 15 | 13 |
|  | Itraconazole | 0.03 | 0.53 | 0.45 | 0.1 | 0.1 | 14 | 14 |
|  | Voriconazole | 0.03 | 0.34 | 0.66 | 0.13 | 0.16 | 18 | 18 |
|  | Posaconazole | 0.03 | 0.5 | 0.5 | 0.08 | 0.07 | 16 | 16 |
|  | Amphotericin B | 0.5 | 0.24 | 0.24 | 0.17 | 0.17 | 9 | 8 |
|  | Caspofungin | 1 | 0.76 | 0.1 | 0.19 | 0.18 | 7 | 6 |

**Table A4.**
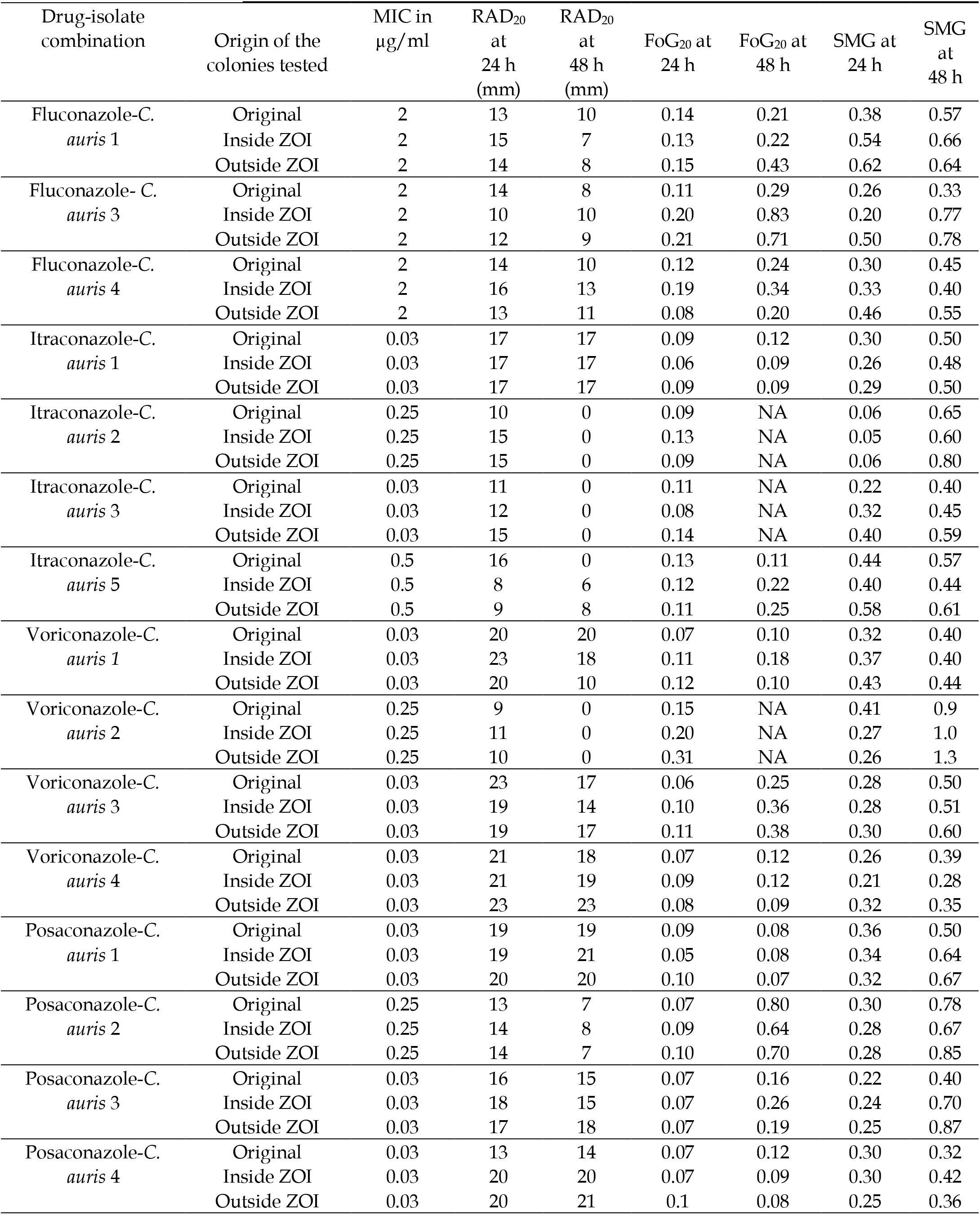
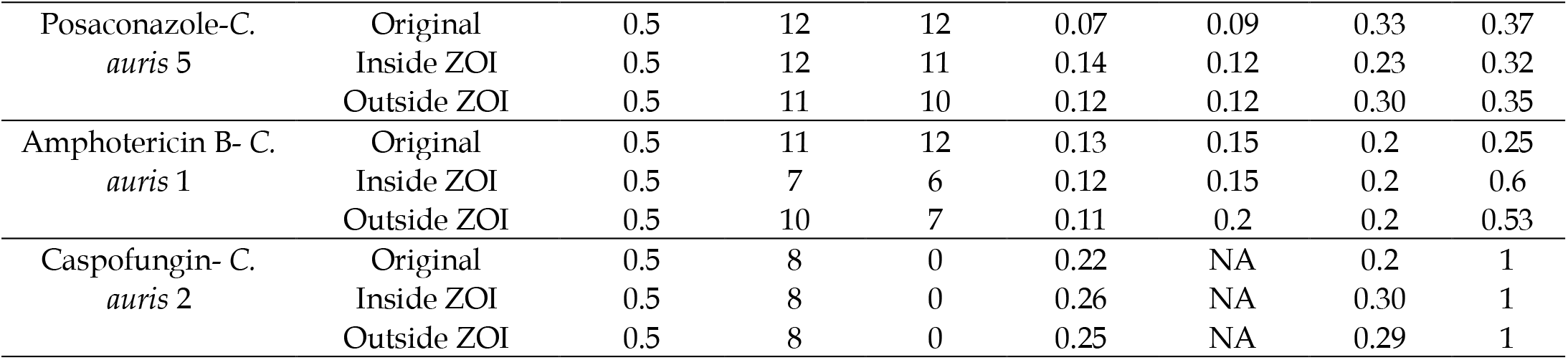
Reversibility of tolerance phenotype among tolerant *Candida auris* isolates against different antifungal agents. Mean radius of the zone of inhibition (RAD), mean fraction of growth (FoG) in the zone of inhibition (ZOI), mean minimum inhibitory concentration (MIC), and mean supra-MIC growth (SMG) values obtained for *C. auris* isolates sub-cultured from inside and outside the ZOI and treated with the azole, polyene, and echinocandin antifungal drugs. MIC, RAD, FoG, and SMG, obtained from *C. auris* colonies isolated from inside and outside the ZOI also did not show any statistically significant differences (Independent t-test, *p* > 0.05 for all values).

**Table A5.** Effect of chloroquine (CLQ) on *Issatchenkia orientalis* and *Candida parapsilosis* reference strains. Mean FoG: Fraction of growth; Mean RAD: Radius of the zone of inhibition. *p*-value was obtained by comparing RAD at 48 hours with and without chloroquine.

|  |  | Without CLQ<br>FoG<br>at 48 hours | With CLQ<br>FoG<br>at 48 hours | Without CLQ<br>RAD (mm)<br>at 48 hours | With CLQ<br>RAD (mm)<br>at 48 hours | Paired<br>t-test |
| --- | --- | --- | --- | --- | --- | --- |
| *C. parapsilosis*<br>ATCC 22019 | Fluconazole | 0.14 | 0.08 | 14 | 9 | $p = 0.195$ |
| | Posaconazole | 0.07 | 0.06 | 17 | 16 | $p = 0.272$ |
| *I. orientalis*<br>ATCC 6258 | Fluconazole | NA | 0.11 | 0 | 7 | $p = 0.032$ |

## Appendix B Supplementary Figures

**Figure B1:**
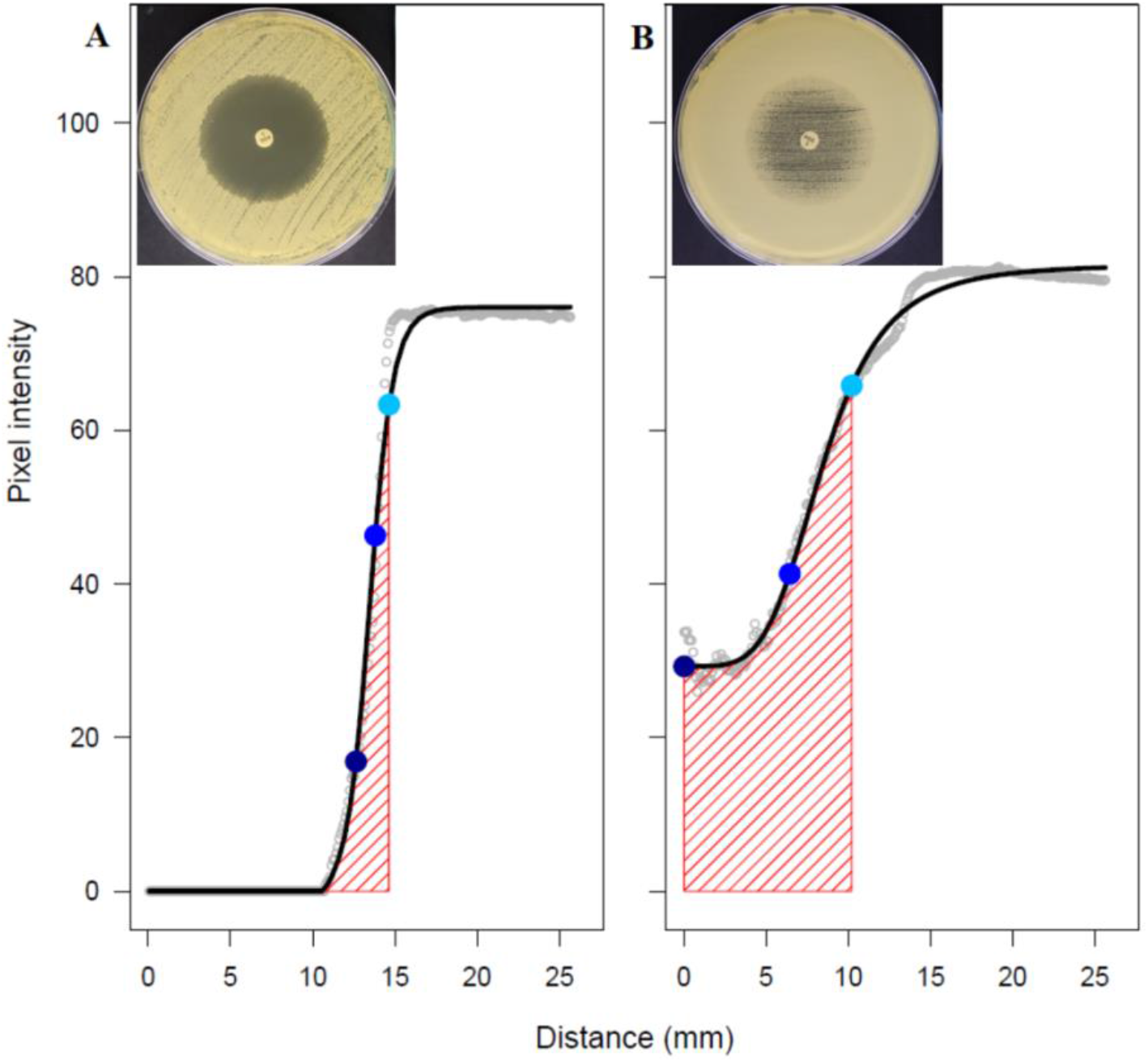
Quantification of antifungal tolerance in a disk diffusion assay using the image analysis program *diskImageR* [29]. Pixel intensity corresponds to the cell density, and its average is measured for 72 radii, every 5 ° from the center of the disk (grey dots). The radius of the zone of inhibition and fraction of growth are measured in three areas where 20 %, 50 % and 80 % of the growth is inhibited (light blue, blue, and dark blue circles, respectively) after **(A)** 24 hours of incubation and **(B)** 48 hours of incubation. The representative data in this figure was obtained from images of a disk diffusion assays for *C. auris* (isolate 2) exposed to posaconazole (insets of (A) and (B)).

**Figure B2.**
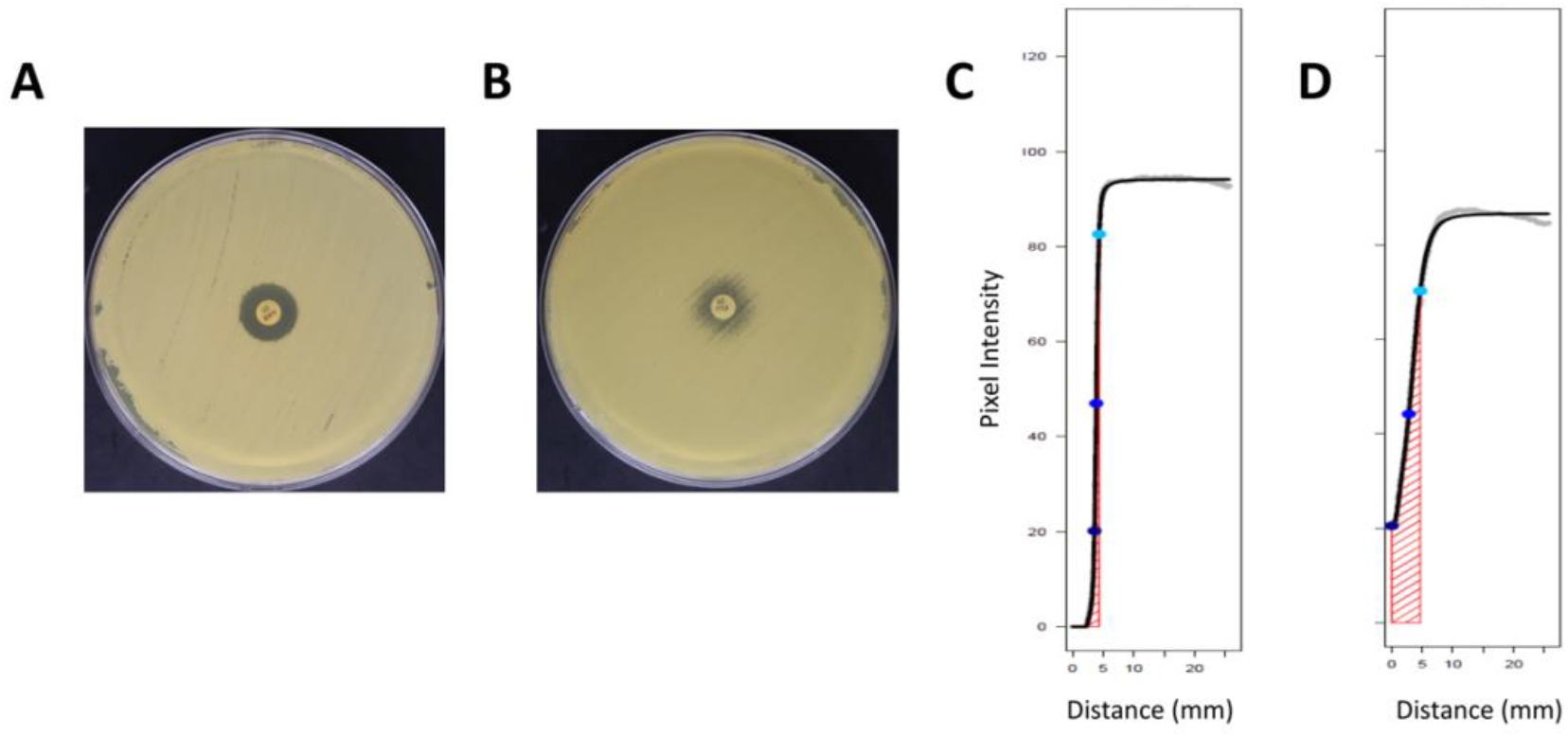
Detecting tolerance in *Candida auris* from disk diffusion assays (DDAs) using *diskImageR*. **(A)** Representative DDA image of *C. auris* (isolate 1) after 24 hours of exposure to amphotericin B (AMB). **(B)** Representative DDA image of *C. auris* (isolate 1) after 24 hours of exposure to fluconazole (FLU). **(C)** Quantification tolerance (shown in the pink zone) from the DDA shown to FLU in (A) using *diskImageR* [29]. **(D)** Quantification tolerance from the DDA shown to AMB in (B) using *diskimageR*.

**Figure B3.**
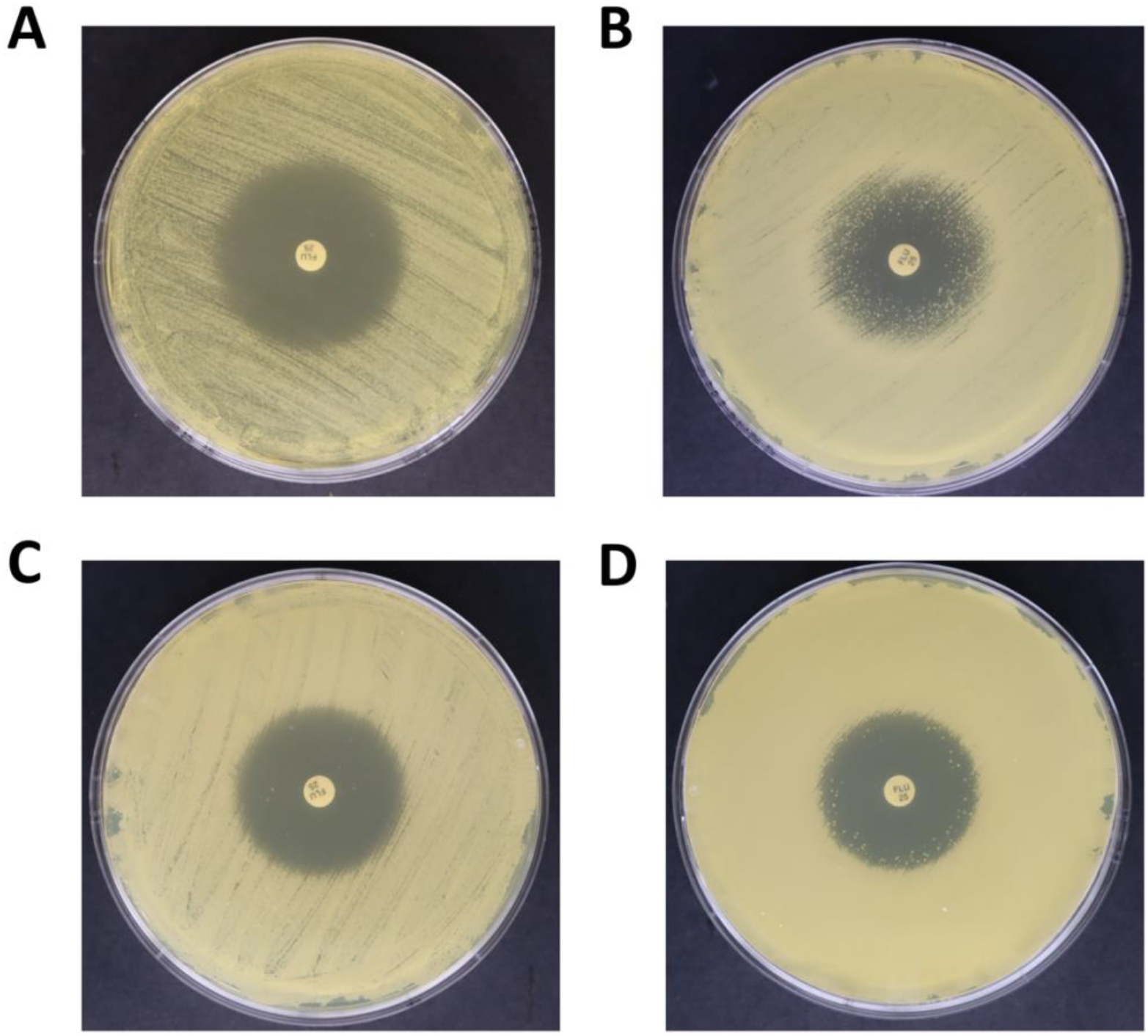
Representative disk diffusion assays (DDA) images of fluconazole (FLU) tolerance in *Candida auris* and *Candida parapsilosis*. **(A)** DDA of *C. auris* (isolate 1) after 24 hours of exposure to FLU. **(B)** DDA of *C. auris* (isolate 1) after 48 hours of exposure to FLU. **(C)** DDA of *C. parapsilosis* after 24 hours of exposure to FLU. **(D)** DDA of *C. parapsilosis* after 24 hours of exposure to FLU.

**Figure B4.**
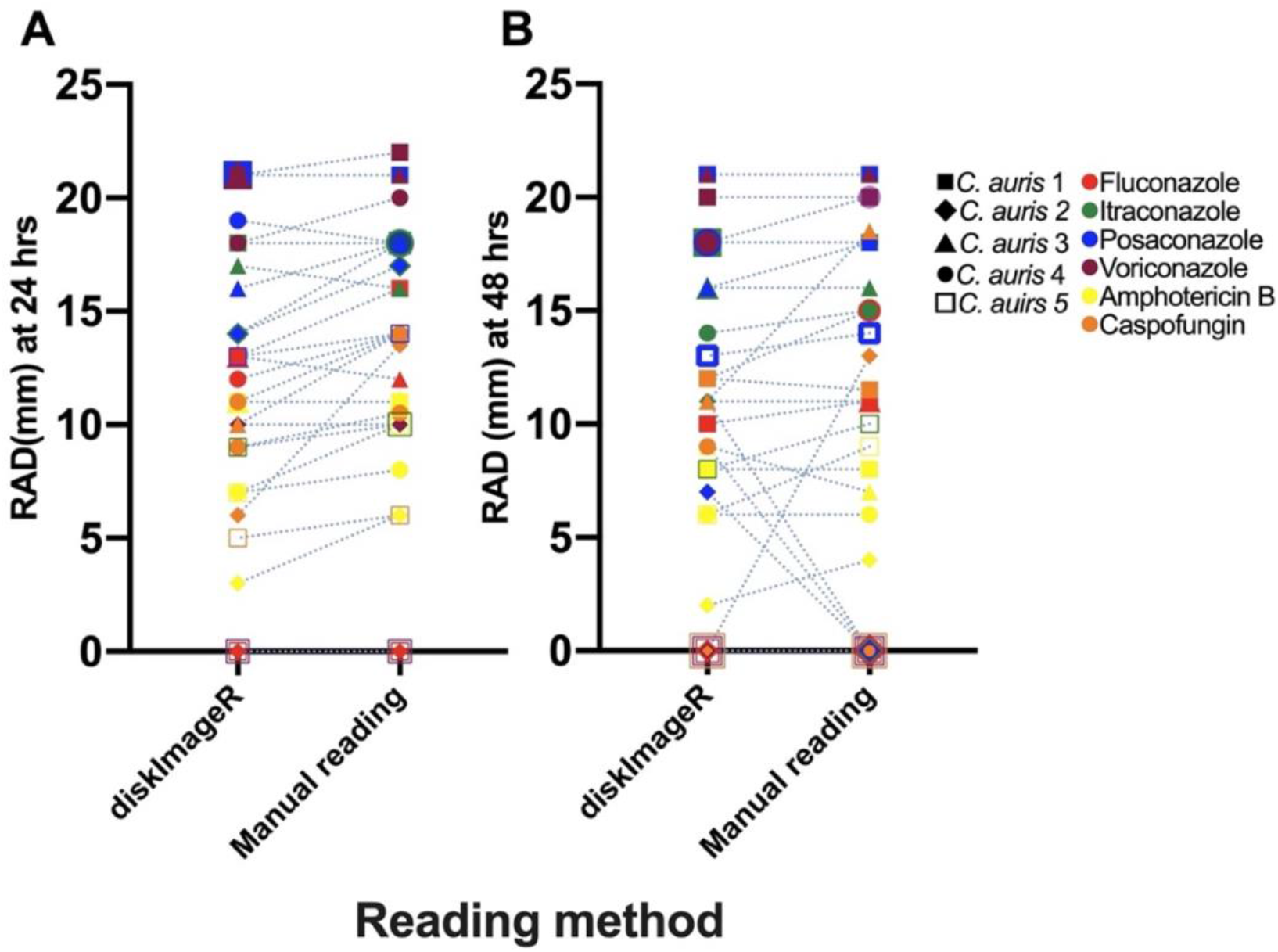
Comparison between *diskImageR* and manual radius of the zone of inhibition (RAD) measurements. **(A)** Mean RAD: Radius of the zone of inhibition measured by *diskImageR* [29] and manually (see Section 2.5) at 24 hours and **(B)** at 48 hours.

**Figure B5.**
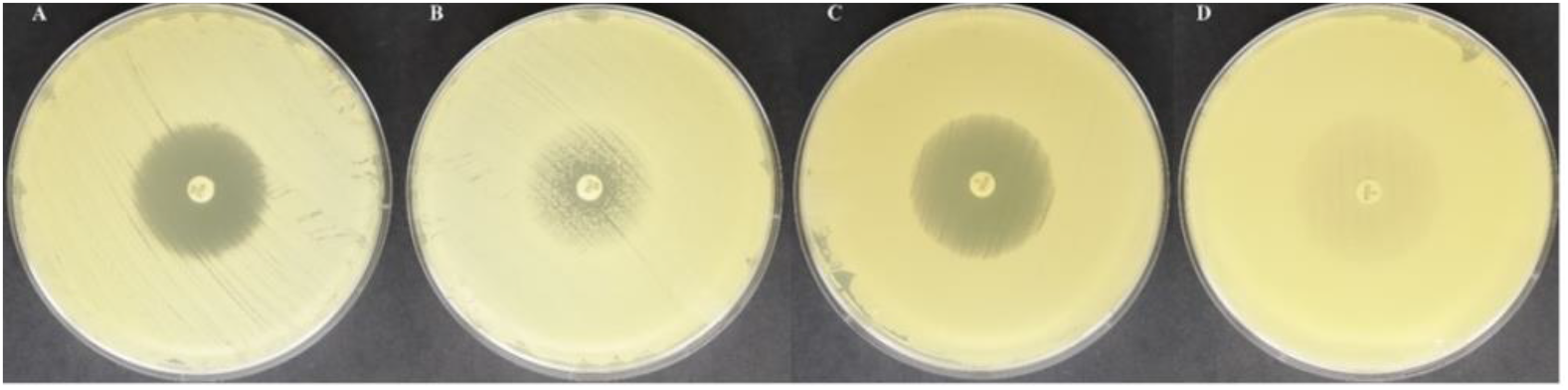
Azole tolerance in *Candida auris*. Disk diffusion assay (DDA) of fluconazole for *C. auris* (isolate 1) after **(A)** 24 hours of growth and **(B)** 48 hours of growth. DDA of posaconazole for *C. auris* isolate 2 after **(C)** 24 hours of growth and **(D)** 48 hours of growth.

**Figure B6.**
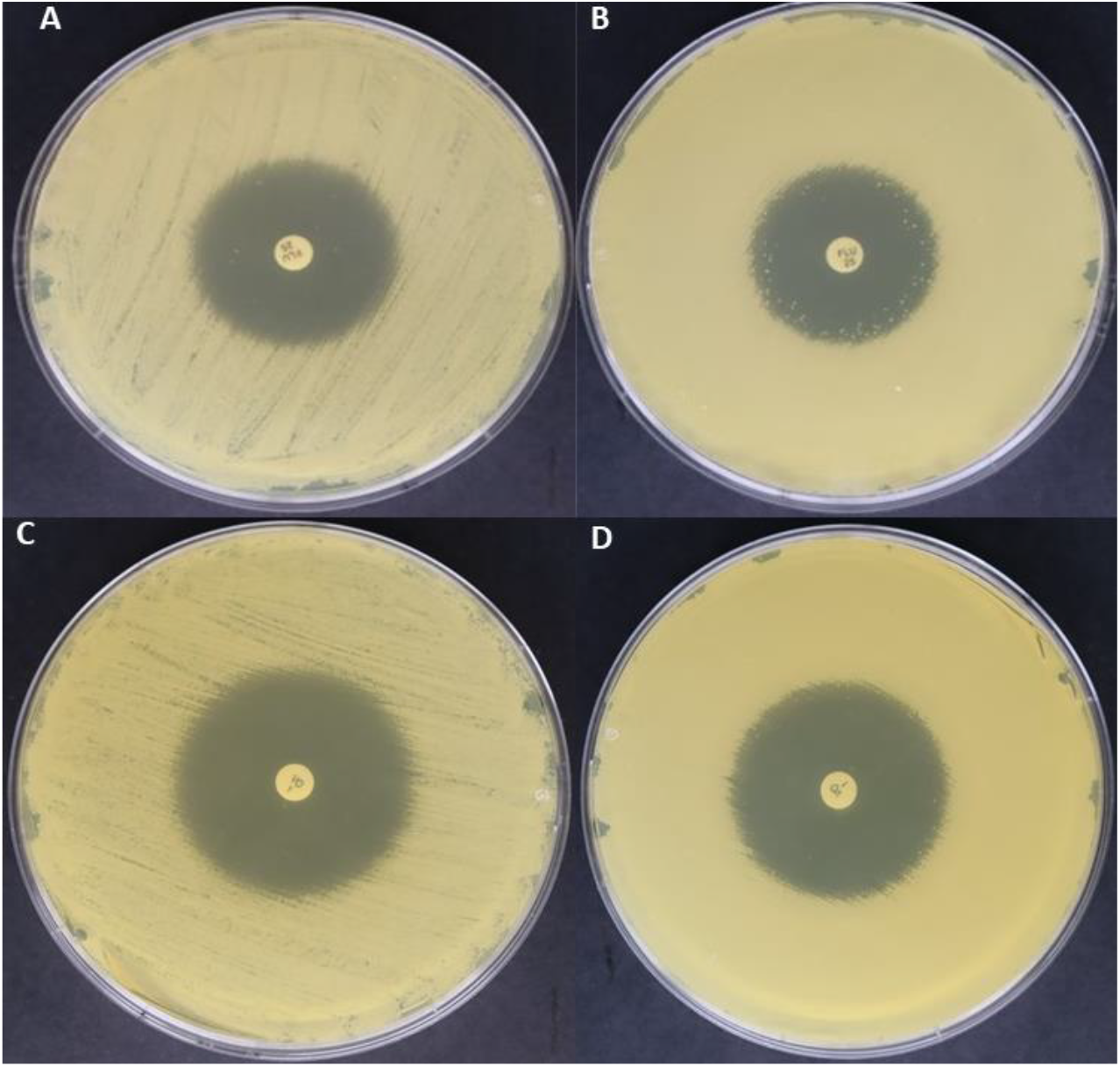
Azole tolerance in *Candida parapsilosis* reference strain. Tolerant *C. parapsilosis* colonies in the zone of inhibition (ZOI) after **(A)** 24 hours and **(B)** 48 hours of fluconazole treatment. Tolerant *C. parapsilosis* colonies in the ZOI after **(C)** 24 hours and **(D)** 48 hours of voriconazole treatment.

**Figure B7.**
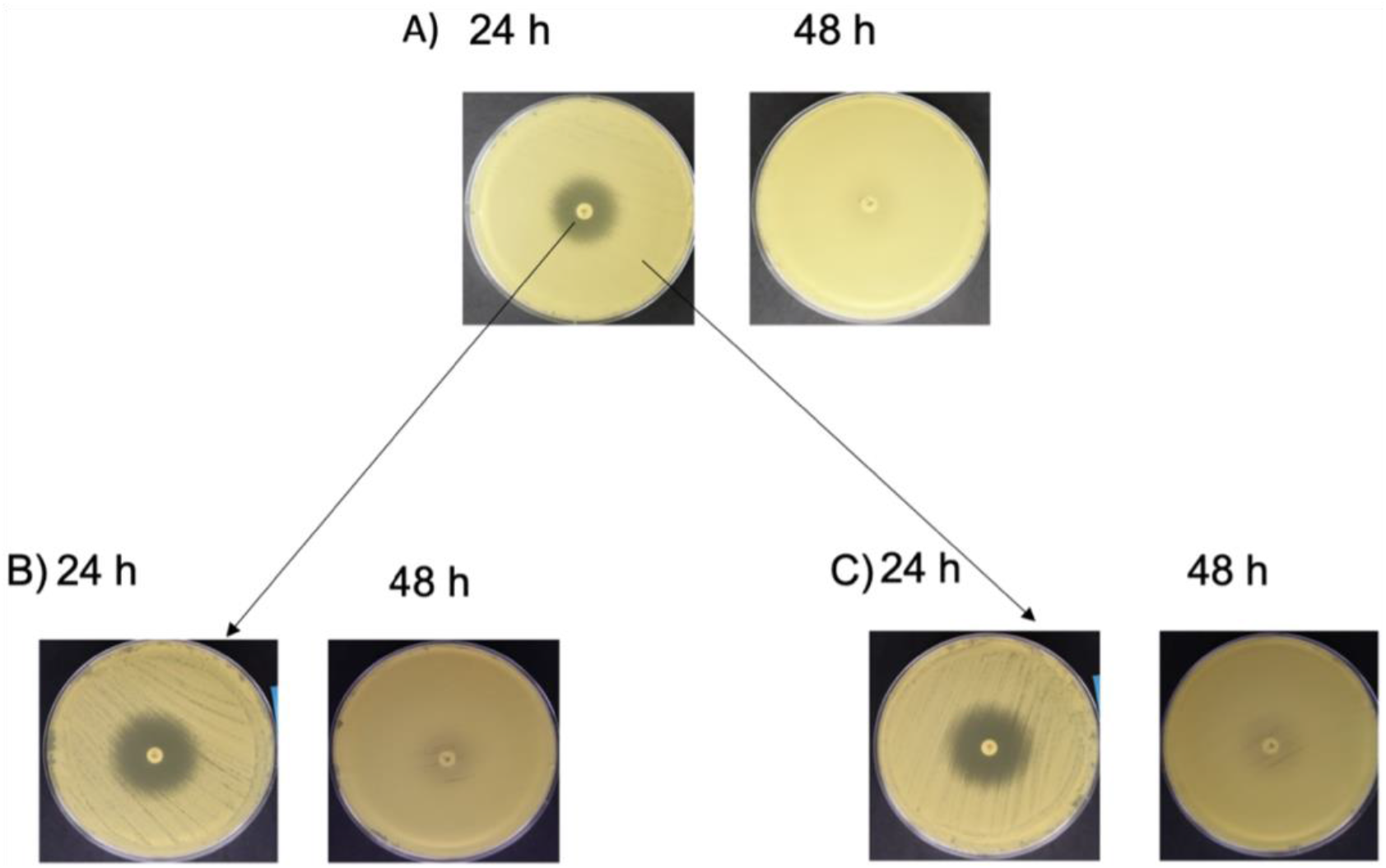
Reversibility of tolerance in a representative *Candida auris* isolate 2 against voriconazole. Disk diffusion assay after 24 and 48 hours for **(A)** *C. auris* original isolate 2, **(B)** colonies isolated and sub-cultured from inside the zone of inhibition (ZOI) of the original plate, and **(C)** colonies isolated and sub-cultured from outside the ZOI of the original plate.

**Figure B8.**
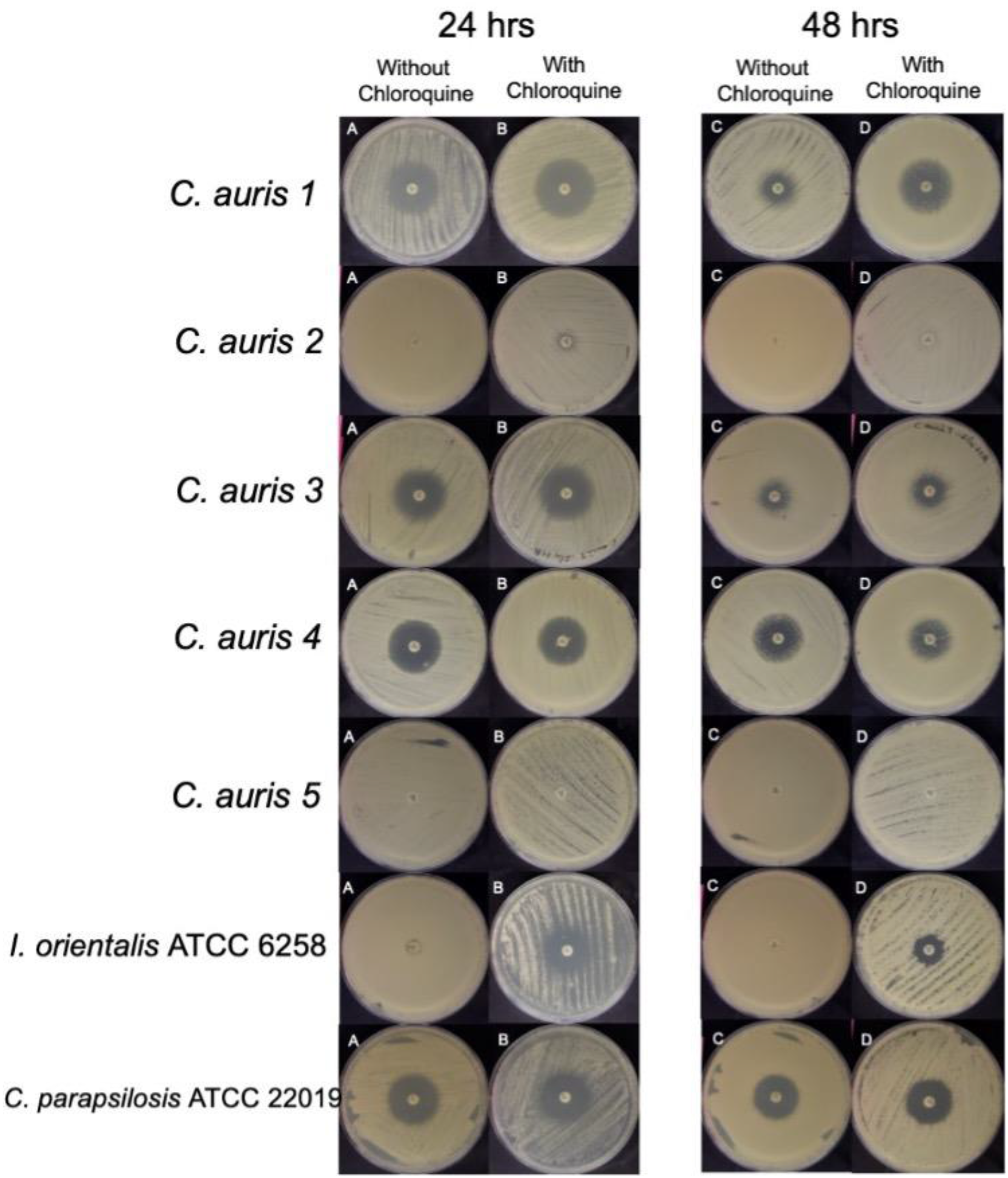
Disk diffusion assays (DDAs) of antifungal-adjuvant treatment in *Candida auris* isolates and *Issatchenkia orientalis* and *Candida parapsilosis* reference strains. DDAs with fluconazole (FLU; 1st column) and with FLU combined with chloroquine (2nd column) against five *C. auris* isolates and two reference strains after 24 hours (left) and 48 hours (right).

**Figure B9.**
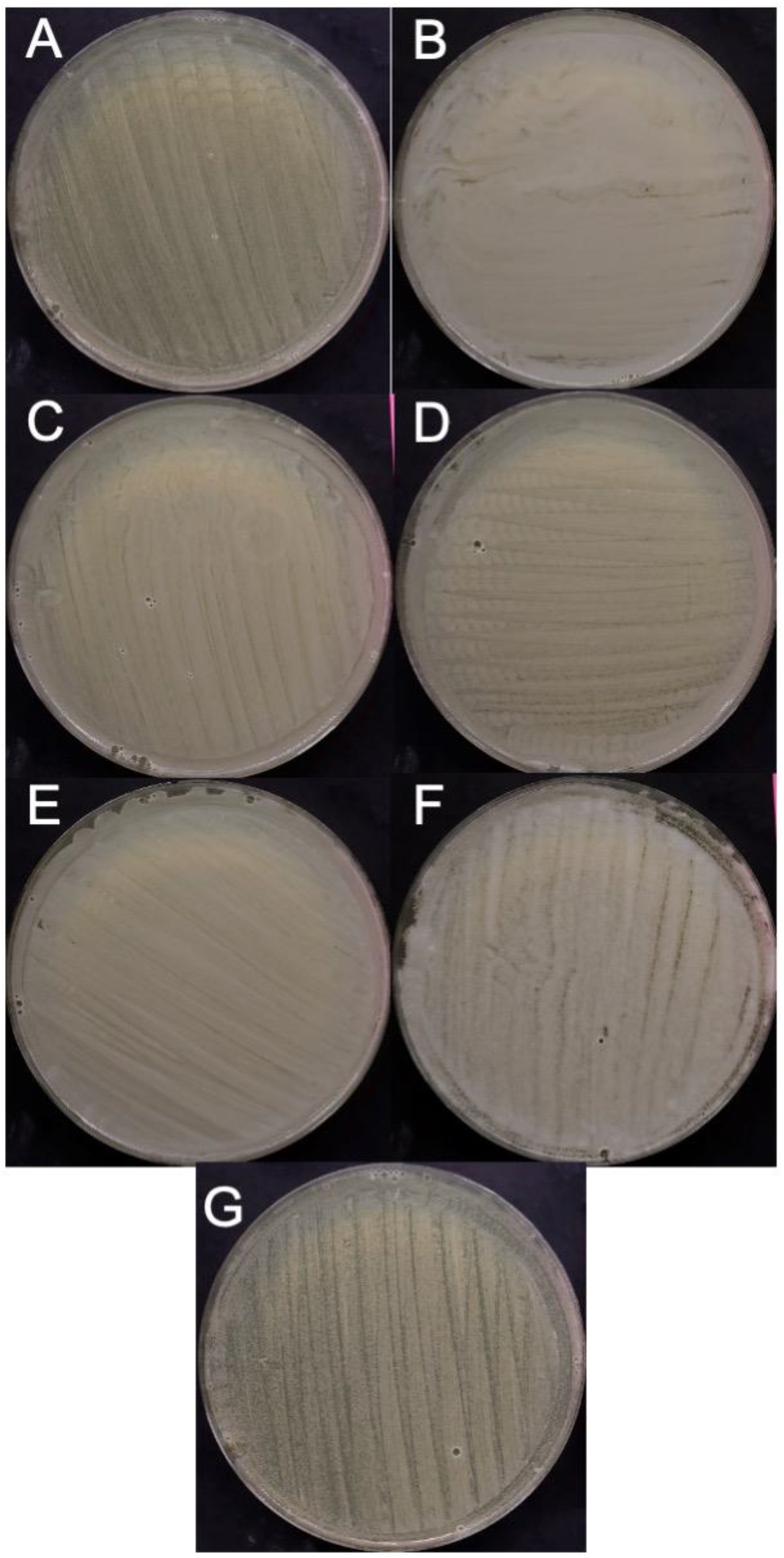
*Candida auris* isolates and *Candida parapsilosis and Issatchenkia orientalis* reference strains growing on Mueller-Hinton agar (MHA) media with chloroquine. Images of *C. auris* isolates 1-5 **(A-E)***, I. orientalis* **(F)**, and *C. parapsilosis* **(G)** grown on MHA plus glucose-methylene blue agar plates with 1,031.8 μg/mL chloroquine diphosphate salt.

